# Long-term acclimation to different stress types: revealing tradeoffs between mesophyll and stomatal conductance

**DOI:** 10.1101/340786

**Authors:** Yotam Zait, Ilana Shtein, Amnon Schwartz

## Abstract

*Ziziphus spina-christi*, a thermophilic tree, became more abundant in the Mediterranean, presumably due to increased winter temperatures. In order to predict the plant acclimation to future climate changes, we attempted to understand which factors underlie photosynthetic stress acclimation.

Stress acclimation to three major long-term abiotic stresses (drought, salinity and temperature) was investigated by measuring growth, gas exchange, chlorophyll fluorescence and leaf structure. We developed a stress index that allowed to precisely define stress level, enabling a comparison between stress types. For each stress, photosynthesis-limiting factors were compared: stomatal conductance (*g_s_*), mesophyll conductance (*g_m_*) and maximum capacity for Rubisco carboxylation (*V_cmax_*).

Photosynthesis under all stresses was limited mostly by *g_s_* and *g_m_* (80-90%); whereas biochemistry (*V_cmax_*) made a minor contribution (10-20%). The relative contribution of *g_s_* and *g_m_* on photosynthetic limitation was influenced by stress type. During acclimation to drought or salinity, photosynthesis was limited by a decline in *g_s_*, while intolerance to low temperatures was driven by decline in *g_m_*. Low mesophyll-imposed limitation was the best predictor of abiotic stress tolerance.

The results demonstrate how warming climate benefits thermophilic species. Moreover, current work gives methodology for stress studies, and defines the main factors underlying the plant response to climate change.

**Highlight:** We have compared the photosynthesis limitation resulting from long-term acclimation to three major a-biotic stresses (drought, salinity and temperature) to understand which limiting-factor are dominant under each type of stress.

## Introduction

An increasing number of studies have shown that climate change affects a wide range of tree species (Saxe et al., 2001; Walther et al., 2002). Climate change can influence habitats in several ways, including increased incidence of extreme temperatures, reduced precipitation and an increase in soil salinity (IPCC, 2014; McDowell et al., 2011). All of these can have a deleterious effect on tree productivity and survival and significantly influence biodiversity (Allen *et al*., 2010). For this reason, it is important to understand how trees acclimate to environmental changes, particularly drought, increased salinity and increased temperature. Predicting stress-acclimation behaviors is complex, since under natural conditions, trees are exposed to various environmental changes that require them to cope with multiple types of stress simultaneously (Niinemets, 2010). Moreover, stress acclimation is a gradual process that involves structural, physiological and biochemical adjustments (Flexas *et al*., 2006a; Galmés *et al*., 2006).

Growth and leaf photosynthetic capacity are the main processes affected by abiotic stress (Chaves *et al*., 2002). Therefore, studies of photosynthesis under stress conditions can help to explain how different species acclimate and adapt to their environments (Flexas *et al*., 2014). It is common practice to consider reductions in photosynthesis as a consequence of three limiting factors (Grassi & Magnani, 2005), specifically decreased CO_2_ concentrations at the carboxylation site caused by limitations on the diffusion of CO_2_ through stomata and mesophyll conductance (stomatal limitation – *L_s_* and mesophyll limitation – *L_m_*) and the capacity for Rubisco carboxylation-*V_cmax_* (biochemical limitations – *L_b_*). Though photosynthetic responses to numerous types of environmental stress have been studied extensively, the majority of quantitative limitation analyses have addressed short-term stress responses (Galmés *et al*., 2007a; Flexas *et al*., 2009; Galle *et al*., 2009, 2011; Pérez-López *et al*., 2012; Chen *et al*., 2015; Wang *et al*., 2017). Short-term responses may not reflect important structural and physiological adjustments that occur under natural conditions (Zhou *et al*., 2016). A few studies have addressed long-term acclimation to drought (Diaz-Espejo *et al*., 2007; Perez-Martin *et al*., 2009; Cano *et al*., 2014; Tomás *et al*., 2014) and growth temperatures (Yamori *et al*., 2006; Warren, 2008; Perdomo *et al*., 2016) but we know of no previous quantitative limitation analyses reports on long-term acclimation to increased soil salinity. Quantitative limitation analyses have also been used to study the stress responses of trees over their seasonal cycles (Grassi and Magnani, 2005; Limousin *et al*., 2010; Misson *et al*., 2010; Egea *et al*., 2011). To date, there are no clear conclusions as to which is the most dominant limiting factor (*L_s_, L_m_* and *L_b_*) for long-term acclimation to stress (Flexas *et al*., 2008). For this reason, it is important to test each type of stress separately, to facilitate comprehensive comparisons. To the best of our knowledge, there have been no previous comparative studies of quantitative limitations on photosynthesis caused by different stress factors in a single species.

*Ziziphus spina-christi* (Rhamnaceae) is a thermophilic tree that is able to survive in arid areas (Saied *et al*., 2008). The species’ natural range of distribution extends from Sudan and Ethiopia to North Africa, India and the Eastern Mediterranean. Its fruits are eatable and traditionally the leaves have a medicinal value and the branches are used as valuable firewood. Over the last few decades, *Ziziphus spina-christi* has become more abundant and more widespread in the Eastern Mediterranean region, presumably as a result of higher winter temperatures (BioGIS, 2018). *Ziziphus spina-christi* is basically evergreen, but under conditions of low temperature in the winter and/or severe drought in the summer, the tree is deciduous or semi-deciduous (i.e., facultative deciduous) (Al Yamani *et al*., 2018). Several questions have been raised regarding changes in the distribution of *Ziziphus spina-christi*: 1. Which environmental parameters define the limits of its growth and photosynthesis? 2. How does the partitioning of photosynthetic limitation (i.e., stomatal, mesophyll, biochemical) differ under different types of stress? 3. Can we use our knowledge of photosynthetic acclimation to stress to predict shifts in this tree’s abundance and natural range of distribution?

To address these questions, under controlled environmental conditions, we thoroughly examined acclimation to three types of environmental stress: drought, salinity and temperature. For each type of stress, we examined four levels of severity. It is important to note that there are various and conflicting definitions of stresses (Levitt, 1980; Lichtenthaler, 1996; Gilbert and Medina, 2016). For this reason, relating photosynthetic limitations directly to the stress severity is a challenge. The types of stress examined in this study were defined in a comparative and precise manner in terms of their impact on the following properties: growth, gas exchange, chlorophyll fluorescence, leaf structure, leaf turgor pressure and leaf hydraulic conductance. Such strict differentiation leads to a better understanding of variations in photosynthesis and to better predictions of plant responses over the course of acclimation to environmental change.

## Materials and Methods

### Plant material

The study was conducted using the thermophilic tree *Ziziphus spina-christi* (Rhamnaceae). One-year-old saplings were obtained from a public nursery (JNF; Gilat, Israel). All saplings originated from uniform plant material. The saplings were kept in a controlled greenhouse before the stress experiments. They were irrigated with a solution that contained the following concentrations of nutrients: 2.0 ± 0.1 mM Ca, 1.2 ± 0.2 mM Mg, 0.27 ± 0.04 mM NH4, 4.6 ± 0.1 mM NO3, 0.30 ± 0.02 mM P and 2.3 ± 0. 2 mM K.

### Stress experiments

#### Drought experiment

All saplings were grown in 10-L containers filled with highly porous organic planting soil inside a greenhouse (30/22°C day/night, VPD=1.8-2.2 kPa). Full-scale lysimeters were used to measure evapotranspiration (ET), to determine the initial required water supply and to induce the desired levels of drought stress. We later monitored the stress levels via daily measurements of volumetric water content (VWC), carried out using a 5TM soil moisture sensor (Decagon Devices) and measurements of leaf water potential (*LWP*) taken using a pressure bomb (model Ari-Mad, Israel). The period of drought lasted 120 days, in order to let the plants acclimate and grow new leaves. Well-watered plants (WW) received 120% ET. For those plants, pot VWC was kept at 40% and *LWP* was kept between −0.5 and −1 MPa. For the mild-stress treatment (WS1), we applied deficit irrigation of 20% ET, which led to a VWC of 22 ± 1.7%. For the high-stress treatment (WS2), plants received only 5% of the initial ET, which led to a VWC of 10.2 ± 2.6%. Severely stressed plants did not receive any water at all and were evaluated after 10 days when VWC had reached zero and *LWP* exceeded −1.8 MPa.

#### Salinity experiment

All saplings were grown in 10-L containers filled with Perlite-2. After 60 days, the plants were subjected to four different irrigation-salinity treatments with NaCl concentrations of 0, 30, 60 and 90 mM. Those treatments were continued for 140 days to allow the plants to acclimate and grow new leaves.

To determine stress levels on a leaf basis, leaf fresh weight (FW) was determined. Leaves were dried in a 70°C oven for 7 days and the leaf Na^+^ concentration was measured using inductively coupled plasma (ICP; ICP-AES, Arcos, Spectro, Kleve, Germany).

#### Temperature experiment

This experiment was conducted in an enclosed greenhouse (phytotron). All saplings were grown in 10-L containers filled with highly porous organic planting soil. In order to allow for gradual acclimation, plants were grown for 40 days at 28/22°C (day/night) before being subjected to the different temperature treatments. Those treatments were: 34/28°C, 28/22°C, 22 /16°C and 16/10°C. Greenhouse VPD was kept in the range of 1.6-1.9 kPa in all treatments. We also used a separate growth room kept at 10/5°C in order to induce severe temperature stress. The experiment lasted 150 days to allow the plants to acclimate and grow new leaves.

### Gas exchange and chlorophyll a fluorescence measurements

Gas-exchange measurements were conducted with the LiCor-6400 portable photosynthesis measurement system, equipped with a 2-cm^2^ fluorescence leaf chamber (6400-40 Leaf-Chamber Fluorometer; LiCor Inc., Lincoln, NE, USA). All measurements were carried out on leaves that had grown under the specific stress conditions. Measurements were taken using the youngest fully expanded leaves (usually between the 5th and 7th from the apical shoot). Measurements were conducted at a saturated light intensity (1500 μmol photons m^−2^ s^−1^), with air flow of 300 μmol air s^−1^ at ambient temperature and RH of 60-50%. Block temperature was set to 32 ^o^C in the drought and salinity experiments. Measurements were carried out between 8:00-11:00 (just before mid-day depression) only on clear days. The diffusional leakage was corrected using the empty chamber method according to Rodeghiero *et al*., (2007). Diffusion coefficients for CO_2_ were K_CO_2__=0.24 for the 2cm^2^ leaf chamber and K_CO_2__=0.31 for the 6cm^2^ leaf chamber.

#### CO_2_ and light-response curves

After a leaf was clamped to the LiCor-6400 chamber, we allowed 15 min of acclimation to a fixed flux of 400 μmol CO_2_ mol^−1^ air for non-stressed plants and 50 μmol mol^−1^ for stressed plants. After the leaf had acclimated, we combined the CO_2_ response curves for gas-exchange (GE) and chlorophyll a fluorescence (CF). Once the first CO_2_ response curves had been calculated, inlet air was replaced with air containing 2% O_2_, the leaves were acclimated for 10 min to a fixed 400 μmol mol^−1^ and an additional CO_2_ response curve was calculated. Next, light-response curves were calculated in ambient (21% O_2_) and low oxygen (2% O_2_).

The final step was to construct CO_2_ response curves with air containing 21% O_2_ at different PPFD levels, according to the Laisk method (Laisk, 1977). For greater accuracy, we used a LiCor-6400XT photosynthesis measurement system with a red-blue light source and a 6-cm^2^ chamber. CO_2_ response curves were constructed for several PPFD levels (400, 200, 100 and 50 μmol m^−2^ s^−1^) at CO_2_ concentrations of 400, 300, 200, 150, 100, 75, 50 and 25 μmol mol^−1^.

### Leaf water relations

After the GE and CF measurements, the measured leaf was cut and inserted into a pressure bomb, in order to measure its water potential (*Ψ_w_*). The same leaf was then immediately transferred into liquid nitrogen. Samples were also put into 0.5-ml Eppendorf tubes and centrifuged to extract the tissue sap. The osmotic potential (*Ψ_s_*) of the extracted sap was determined using a vapor pressure osmometer (Model 5600, ELITech Group, Puteaux, France). Leaf turgor pressure (*Ψ_p_*) was later calculated using the water potential equation: *Ψ_p_* = *Ψ_w_* – *Ψ_s_*.

To estimate leaf hydraulic conductance, we measured midday stem water potentials (*Ψ_stem_*) by covering leaves, approximately 3 h prior to cutting, with a bag that was impermeable to water vapor. Leaf hydraulic conductance (*K_leaf_*) was estimated using the evaporative flux method (Sack *et al*., 2002), that is, *K_leaf_* = *E*/(*Ψ_stem_* – *Ψ_leaf_*) where E is the transpiration rate measured earlier with the LiCor-6400.

### Analysis of leaf starch and soluble sugars

One hundred mg of fine powder were extracted three times in hot 80% ethanol. Total soluble sugar content was determined using the Anthrone reagent method (Dubois *et al*., 1956). To determine starch content, the dried residue was analyzed following amyloglucosidase digestion.

### Estimation of mesophyll conductance and theoretical considerations

To estimate mesophyll conductance (*g_m_*), we used the variable J (Harley *et al*., 1992) and curve-fitting methods (Ethier and Livingston, 2004). These two methods rely on different assumptions, but showed good agreement (Fig. S2). *g_m_* estimation by the variable J method is expressed as:

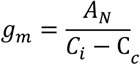

where A_N_ is the net photosynthesis rate and *C_i_* is the CO_2_ concentration inside leaf intercellular airspaces (both determined from GE measurements). *C_c_* is the CO_2_ concentration in the chloroplast stroma, estimated as:

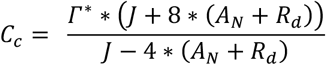

where *J* is electron-transport rate calculated based on chlorophyll fluorescence, calibrated by the GE data (as described below). *R_d_* is the non-photorespiratory respiration in light and *I*^*^ is the apparent photo-compensation point estimated according to the Laisk method (Laisk 1977). Laisk CO_2_ curves were fitted curvilinearly (Tholen *et al*., 2012) and *y* and *x* values of the average of four CO_2_ response intersections were taken as *R_d_* and 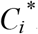. According to von Caemmerer *et al*., (1994), *Γ*^*^ is dependent on *g_m_* and *R_d_*, which are related to one another as follows:

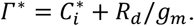

However, since no *gm* value was available prior to the measurements, conversion was not completed and 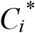 was taken as a good representation of *Γ*^*^ (Qiu *et al*., 2017). To calculate the rate of electron transport (*J*), we constructed CO_2_ and light-response curves in the absence of photorespiration (2% O_2_ in air).

The actual quantum efficiency of photosystem II (*ΦPS*_2_) was calculated according to Genty *et al*. (1989):

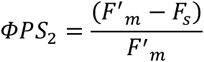

where *F_m′_* is the maximum fluorescence after a light-saturating pulse of ∼8000 μmol photons m^−2^ s^−1^ and *F_s_* is the steady-state fluorescence under continuous light exposure.

The electron transport rate (*J_f_*) was calculated as:

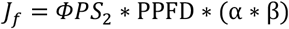

where *α* is the coefficient of leaf absorbance and β is the fraction of absorbed quanta that reach PS_2_.

The quantum yield of CO_2_ fixation (*ΦCO*_2_) was calculated from the GE measurements as: *ΦCO*_2_ =(*A_N_*+*R_d_*)/*PPFD*.

Agreement between GE and CF was predicted on the assumption that under non-(photo)respiratory conditions, there is a linear relationship between *ΦPS*_2_ and *ΦCO*_2_, since CO_2_ fixation is the only sink for electrons (Valentini *et al*., 1995). The theoretical model and experimental observations were limited to the linear region of *ΦCO*_2_ < 0.05, *and ΦPS*_2_ < 0.5. The slope of the linear regression (*k*) and the y-axis intercept (*b*) were used to recalculate the (*α*^*^*β*) for the electron-transport rate (*J_cal_*) based on CF measured under ambient oxygen conditions (O_2_ = 21%) as:

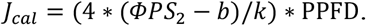

### Considerations for the estimation of *g_m_* under stress conditions

We estimated *g_m_* for stressed plants by allowing the plants to acclimate to low concentrations of CO_2_ (Centritto *et al*., 2003). This method reduces stomatal limitation and maintains sufficient *C_i_* to avoid the negative effect of (photo)respiration on *g_m_* (see Tholen et al., 2012; Théroux-Rancourt et al., 2014). In addition, in severely stressed plants, the epidermis transfers significantly more water vapor than CO_2_, which may bias the estimation of *C_i_* (Boyer *et al*., 1997). Taking all of this into consideration, we measured cuticular conductance (*g_cut_*) for severely stressed leaves by covering the abaxial epidermis with paraffin. Cuticular conductance (*g_cut_*) was found to be 0.005 ± 0.001 mol m^−2^ s^−1^. Measurements of fluorescence quantum yield at several locations on stressed leaves revealed that clear stomatal patchiness could be observed only when stomatal conductance of water vapor (*g_sw_*) was less than 0.05 mol H_2_O m^−2^ s^−1^ (or g_*sc*_>0.0312 mol H_2_O m^−2^ s^−1^). Therefore, in order to avoid any negative effects of stomatal patchiness and dominant cuticular conductance on the estimation of *C_i_*, we decided not to perform any measurements on plants with *g_sw_* lower than 0.05 mol H_2_O m^−2^ s^−1^ (i.e., *g_cut_* that was less than 10% of *g_sw_*). Since the dependence of *g_m_* on *C_i_* is sensitive to measurement error (Pons *et al*., 2009; Gilbert et al, 2011), *g_m_* values were considered reliable only when *dC_c_*/*dA_N_* values were within the criterial range of 10 < *dC_c_*/*dA_N_* < 50, as described in detail by Harley *et al*. (1992).

### Quantitative limitations of photosynthesis

The *A_N_*/*C_i_* curves were converted into *A_N_*/*C_c_* curves and analyzed using the fitting tool developed by Sharkey *et al*. (2007). The maximum carboxylation rate (*V_cmax_*) and rate of photosynthetic electron transport (*J_max_*) were computed. Parameters were adjusted to 25°C to enable comparisons (Bernacchi *et al*., 2001). We later used the *A_N_*/*C_c_* curves to calculate the absolute limitations on net photosynthesis, as described by Grassi and Magnani (2005). In brief and as described in detail by Farquhar *et al*. (1980), light-saturated photosynthesis can be expressed as:

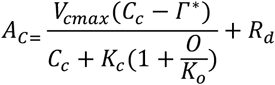

where *A_c_* is the rate of photosynthesis during the Rubisco carboxylation-limited stage, O is the O_2_ partial pressure at the site of carboxylation and *K_c_* and *K_o_* are the Michaelis-Menten constants of Rubisco for carboxylation and oxygenation were taken from Bernacchi *et al*., 2002. The rate of photosynthesis can be limited by substance availability (*Cc*) caused by diffusional restrictions (*g_s_* and *g_m_*) or by biochemical factors (changes in *V_cmax_*). Therefore, total photosynthetic limitations are divided into three components: stomatal limitation (*L_s_*), mesophyll limitation (*L_m_*) and biochemical limitation (*L_b_*). Changes in light-saturated photosynthesis (*ΔA_c_*/*A_c_*) can be expressed in parallel terms of relative change in *g_sc_* (*g_sw_*/1.6), *g_m_* and 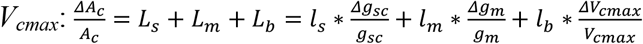

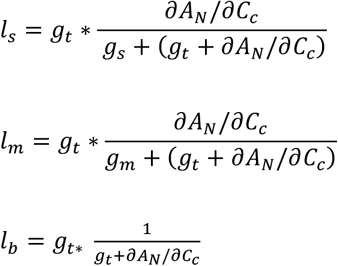

where *g_t_* is the total CO_2_ leaf conductance, calculated as:

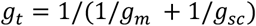

and *∂A_N_*/*∂C_c_* is the first derivative of *A_N_*/*C_c_* curves calculated as:

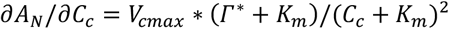

where *K_m_* is the effective Michaelis–Menten constant for CO_2_. Total limitations on the rate of photosynthesis were calculated based on the maximum reference photosynthesis (*A_max_*): *ΔA_N_*/*A_N_* = (*A_max_* – *A_n_*)/*A_N_*.

### Leaf structural analyses

#### Leaf mass per unit area (LMA)

Leaf area was measured using an area meter (model LI-3100, Li-Cor, Lincoln, NE, USA). Leaves were dried in a 70°C oven for 7 days, and then dry weight was measured.

#### Growth measurements

Measurements of the elongation of the main stem were carried out by attaching plastic markers on the apical meristem at a period of every 5 days.

#### Histology

The central area of the uppermost fully expanded leaf was measured. Samples were immediately fixed in FAA (5:5:90, formalin: acetic acid:70% ethanol), dehydrated in a graded alcohol series and embedded in paraffin wax (Paraplast Plus, Leica). Cross-sections were cut 12-μm thick using a rotary microtome (Leica, Germany) and stained in Toluidine Blue O. Samples were viewed and micro-graphed using an inverted microscope imaging system (EVOSTM XL Core, Fisher Scientific).

Image analysis was done using ImageJ software (Rasband, W.S., ImageJ, U. S. National Institute of Health, Bethesda, MD, USA, http://imagej.nih.gov/ij/, 1997–2015). Leaf thickness was measured at three different locations on the cross section. The mesophyll surface area exposed to intercellular airspace per unit leaf area (*S_mes_*/*S*) was calculated according to Syvertsen *et al*. (1995) as *S_m_* = (*L_m_*/*W*) \**F*, where *L_mes_* is the total length of mesophyll cells facing the intercellular air space, *W* is the section width and *F* is the cell-shape curvature correction factor for ellipsoids (Evans et al. 1994). The fraction of the intercellular air space (%*IAS* or *fIAS*) was calculated as 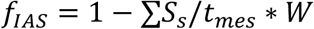, where Σ*S_s_* is the sum of the cross-sectional areas of the mesophyll cells and t_mes_ is the thickness of the mesophyll between two epidermal layers. Stomatal density was measured from epidermal peels, as described by Shtein *et al*. (2011).

### Stress index

The stress index was calculated as a percentage of the difference in physiological variables between the optimal reference (*o*) and the stress value (*s*) using the following equation:

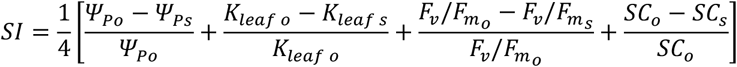

where *Ψ_P_* is the leaf turgor pressure and *K_leaf_* is the leaf hydraulic conductance. *Ψ_P_* and *K_leaf_* are both indicative of leaf water status. *F_v_*/*F_m_* is the dark-adapted chlorophyll fluorescence, indicating the potential efficiency of *PS*_2_. SC is the soluble carbohydrate content, which is indicative of the leaf’s metabolic state.

### Statistical analysis

We used five to seven trees as biological repeats for each treatment and control. We performed three technical repeats for each biological repeat. Analysis of variance (ANOVA) was used (JMP 14 software, SAS Institute Inc., Cary, NC, USA) to identify any statistical differences between treatments. The Tukey-Kramer post hoc test was used to compare the treatments found to be significantly different from one another. Data-fitting was carried out using MATLAB software (MathWorks Inc., Natick, MA, USA).

## Results

### Precise definition of stress

This study focused on three different types of abiotic stress (drought, temperature and salinity) and examined four levels of severity for each type of stress. In order to physiologically define each type of stress and set stress thresholds, measurements of main stem elongation were carried out for all of the stress treatments. The moderate stress threshold was defined as a 50% decline compare to the average of the control growth rate. In the drought-stress experiment, a 50% decline in the growth rate was observed as growth of 0.6 ± 0.06 cm per day (Fig. 1A), which was within the leaf water potential (*LWP*) range of −1 > *LWP* > −1.4 MPa. The lowest *LWP* values that allowed for any growth were between −1.4 and −1.8 MPa. Under those conditions, the growth rate was 0.34 ± 0.03 cm per day, which represented a 72% reduction relative to the control. *LWP* values below −1.8 MPa caused a cessation of growth and were, therefore, defined as severe water stress.

**Fig. 1.**
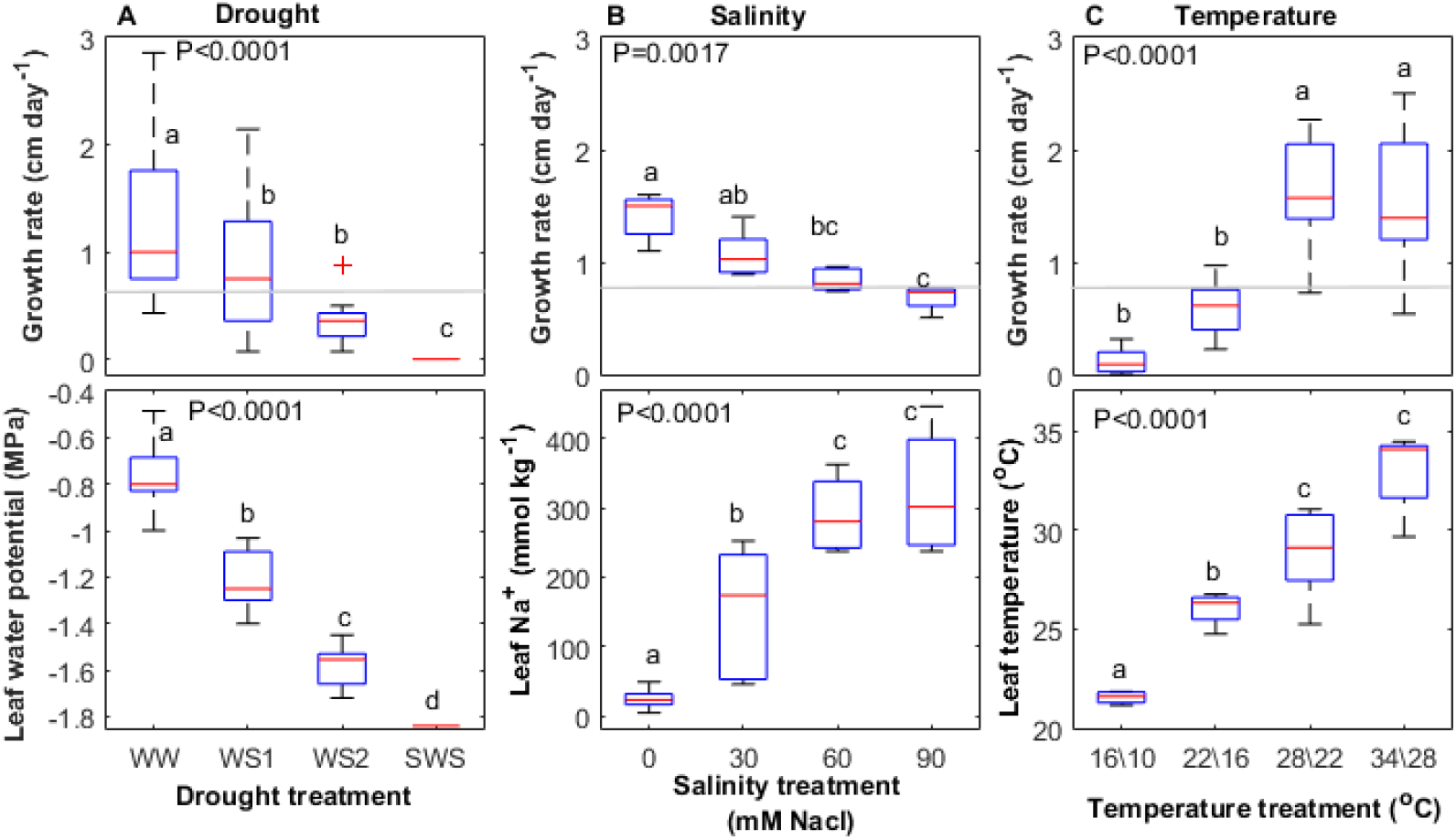
Physiological definition of the types of stress examined, as demonstrated in terms of changes in the growth rate of the main stem and aspects of the leaf stress response. **(a)** The drought experiment included four different irrigation treatments defined by ranges of leaf water potential, **(b)** the salinity experiments were defined in terms of different leaf Na^+^ concentrations and **(c)** the temperature experiment was based on the use of different leaf temperatures. The gray line represents a 50% reduction in the growth rate for each type of stress.

Under salinity stress, a 50% reduction in the growth rate was defined as growth of 0.7 ± 0.05 cm per day, which was observed at a NaCl concentration of 90 mM (Fig. 1B). Therefore, severe salt stress was not reached. For temperature stress, a 50% reduction was defined as growth of 0.77 ± 0.1 cm per day in the 22/16°C treatment, which corresponded to leaf temperatures below 25°C (Fig. 1C). In the 16/10°C temperature treatment, the growth rate was 0.12 ± 0.03 cm per day, which represented a 92% reduction relative to the control.

In order to define GE under various stress conditions, we analyzed the different types of stress in terms of their effects on leaf properties during the acclimation process: leaf water potential for the drought experiment, leaf Na^+^ concentration for the salinity experiment and leaf temperature for the temperature experiment. As shown in Figure 2, the net photosynthetic rate (*A_N_*) declined hyperbolically (*R*^2^ = 0.79, P<0.0001) with the decrease in *LWP* (Fig. 2A), declined nearly exponentially with the increase in leaf Na^+^ (*R*^2^ = 0.66, P<0.0001) (Fig. 2C) and decrease in leaf temperature (*R*^2^ = 0.79, P<0.0001) (Fig. 2E). The maximum capacity for Rubisco carboxylation (*V_cmax_*) exhibited threshold-like behavior in its response to drought and dropped only when *LWP* exceeded −1.5 MPa (Fig. 2B). However, it decreased exponentially with the increase in leaf Na^+^ (*R*^2^ =0.55, P=0.0004) (Fig. 2D) and with the reduction in leaf temperature (*R*^2^ = 0.78, P<0.0001) (Fig. 2F).

**Fig. 2.**
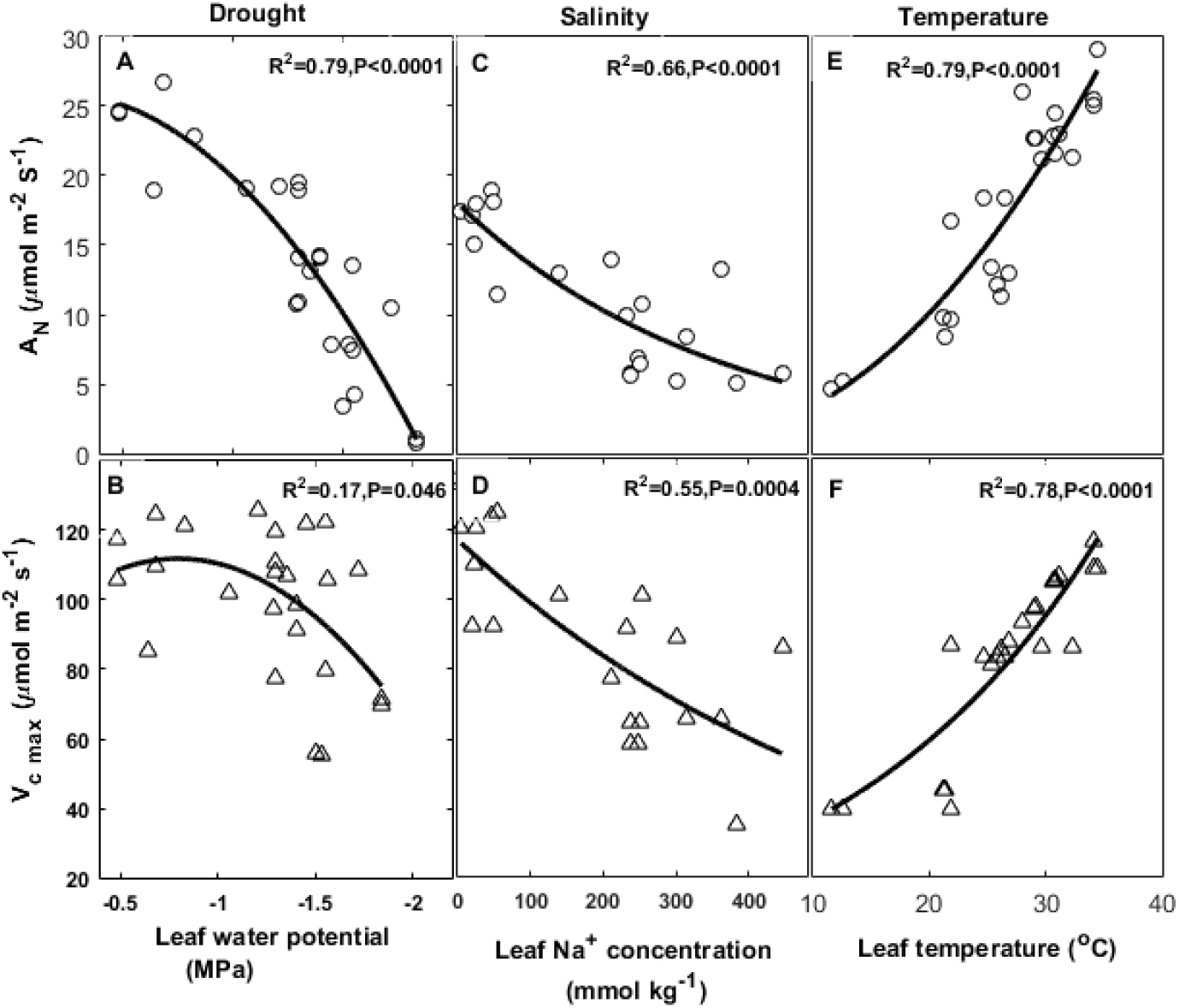
Effects of different types of environmental stress on the rate of photosynthesis (*A_N_*) and the maximum capacity for Rubisco carboxylation (*V_cmax_*). Leaf-level parameters were quantified differently following long-term stress acclimation: **(a)** drought stress was quantified in terms of leaf water potential, **(b)** salinity stress was quantified in terms of leaf Na^+^ concentration and **(c)** temperature stress as quantified in terms of leaf temperature. Continuous lines represent best-fit adjustments.

### Mesophyll to stomatal conductance ratio (*g_m_*:*g_s_*) as a driver of intrinsic water-use efficiency

Under favorable conditions, in all experiments, stomatal conductance to CO_2_ (*g_sc_*) was ∼10-50% higher than *g_m_* (Fig. 3). *g_sc_* and *g_m_* were not coordinated during acclimation to drought and salinity. In the drought experiment, while *g_sc_* decreased exponentially with the decline in *LWP* (*R*^2^ = 0.79), *g_m_* was nearly constant (∼0.16 mol m^−2^ s^−1^) under moderate water stress (-0.5 > *LWP* > −1.5 MPa,) and dropped significantly when *LWP* exceeded −1.5 MPa (Fig. 3A).

**Fig. 3.**
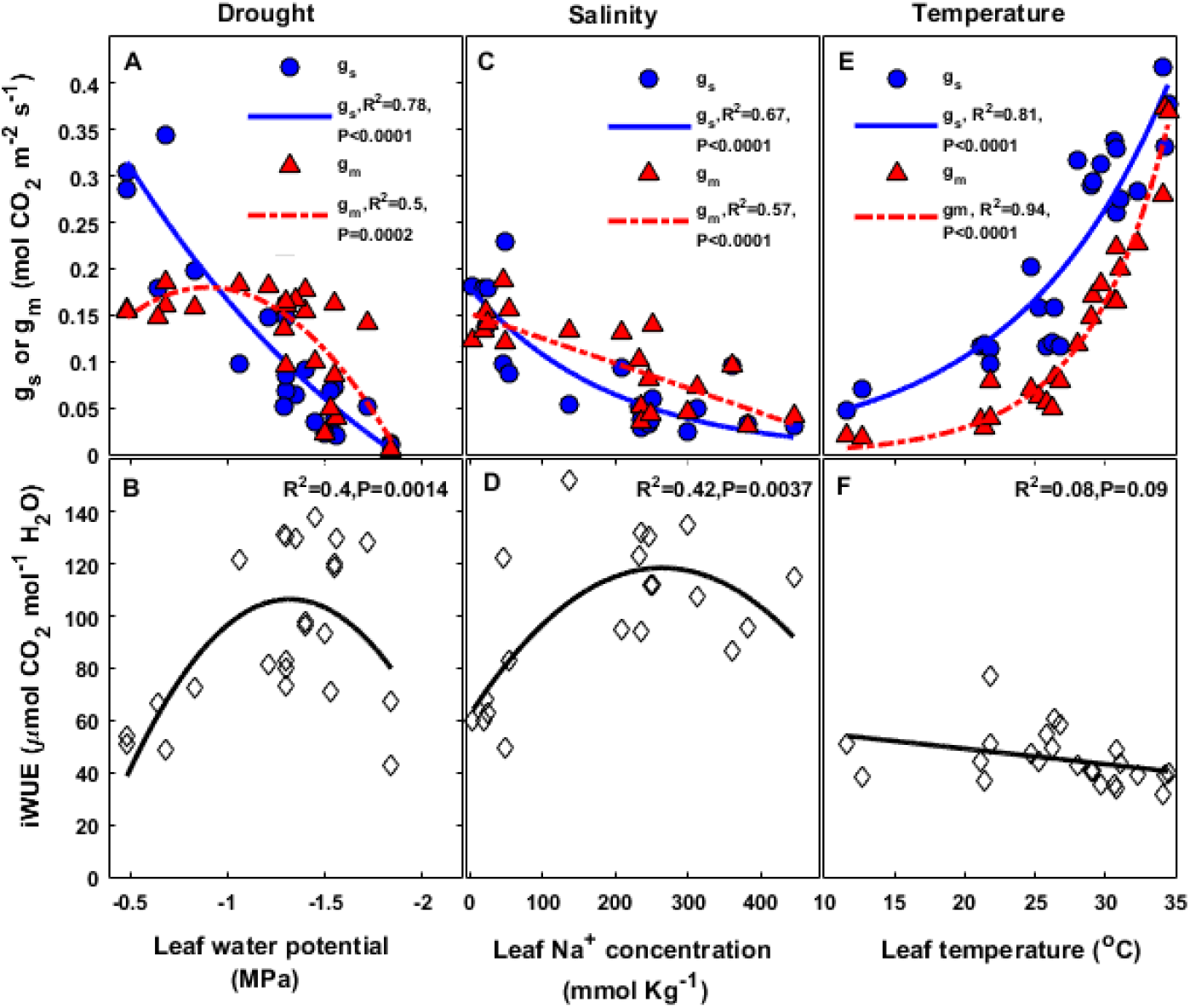
Mesophyll and stomatal CO_2_ conductance (*g_m_* and *g_s_*, respectively) and the change in the leaf-level intrinsic water-use efficiency (*iWUE*) following long-term stress acclimation. Mesophyll conductance and stomatal conductance trends are distinguished by the leaf-level response: **(a)** drought stress as quantified in terms of leaf water potential, **(b)** salinity stress as quantified in terms of leaf Na^+^ concentration and **(c)** temperature stress in terms of leaf temperature. Continuous lines represent the best-fit adjustment for stomatal conductance (blue circles) and broken lines represent the best-fit for mesophyllic conductance (red triangles).

Variation in *g_m_* was relatively high within the range of −1.4 > *LWP* > −1.8 MPa, to the extent that in some plants it was reduced to nearly zero (∼0.05 mol m^−2^ s^−1^); whereas in other plants, it remained high (∼0.15-0.2 mol m^−2^ s^−1^). *g_m_* reached zero under severe water stress (*LWP* < −1.8 MPa). A greater decline in *g_sc_* relative to *g_m_* had a strong impact on intrinsic water-use efficiency (*iWUE*), that is, the ratio of *a_N_* to *g_sw_* (Fig. 3B). There was a moderate increase in *iWUE* between *LWP* values of −0.5 to −1.5 MPa, from ∼40 up to ∼130 μmol mol^−1^, but *iWUE* fell back to its initial lower values when *LWP* was less than −1.5 MPa.

In the salinity experiment (Fig. 3C), *g_sc_* decreased exponentially as leaf Na^+^ increased (*R*^2^ = 0.67, P<0.0001), while *g_m_* decreased moderately and linearly (*R*^2^ = 0.57, P<0.0001). *g_sc_* was greater than *g_m_* (∼0.18 vs. 0.15 mol m^−2^ s^−1^) at low leaf Na^+^ levels (< 50 mmol kg^−1^). When Na^+^ levels were raised to above 130 mol kg^−1^, *g_m_* was larger than *g_sc_* (∼0.05 vs. 0.1 mol m^−2^ s^−1^). As in the case of drought, when there was a sharp decline in *g_sc_* relative to *g_m_*, within the range of 0 < leaf Na^+^ < 200 mmol kg^−1^, we observed an increase in *iWUE* from to 60 to 120 μmol CO_2_ mol^−1^ H_2_O (Fig. 3D).

In the temperature experiment (Fig. 3E), *g_sc_* and *g_m_* both decreased exponentially (*g_sc_*: R^2^=0.81, P<0.0001, *g_m_*: R^2^=0.94, P<0.0001) over the course of acclimation to decrease in leaf temperatures. However, *gm* decreased more sharply and *gm* values were lower than *gsc* values across all of the examined leaf temperatures, which led to low (∼40-50 μmol CO_2_ mol^−1^ H_2_O) and relatively constant *iWUE* across the entire range of examined leaf temperatures (Fig. 3F).

### Limitation of photosynthesis during long-term acclimation to stress

As shown in Figure 4, across all of the examined types of stress, the net photosynthesis rate (*A_N_*) was limited mostly by factors related to the diffusion of CO_2_: stomatal limitation and mesophyll limitation (*L_s_* and *L_m_*). In spite of the obvious decline in *V_cmax_*, biochemical limitation (*L_b_*) of the rate of photosynthesis remained relatively low for all types of stress (∼5% drought ∼15% salinity ∼20% temperature). In the drought experiment (Fig. 4A), the total photosynthesis limitation (*L_t_*) increased exponentially (*R*^2^ = 0.72, P<0.0001) as *LWP* decreased. *L_s_* was hyperbolically correlated with the ongoing decrease in *LWP* (*R*^2^ = 0.51, P<0.0001). *L_m_* was relatively constant (∼15%) under well-watered and moderate drought conditions (−0.5 > *LWP* > −1.5 MPa) and increased sharply to ∼75% when *LWP* was above - 1.5 MPa (*R*^2^ = 0.74, P<0.0001). In the salinity experiment (Fig. 4B), *L_t_* increased hyperbolically (*R*^2^ = 0.79, P<0.0001), reaching a plateau (∼70%) at a leaf Na^+^ concentration of ∼ 200 mmol kg^−1^. *L_s_* was the most dominant limiting factor along all of the examined levels of leaf Na^+^; it increased hyperbolically (*R*^2^ = 0.54, P=0.0005) and plateaued (∼40%) when leaf Na^+^ was ∼200 mmol kg^−1^. *L_m_* exhibited more moderate behavior, as it was ∼10% at low leaf Na^+^ (<200 mmol kg^−1^), and increased inversely to *L_s_* (*R*^2^ = 0.41, P=0.0043), up to ∼30-40%, only when leaf Na^+^ exceeded 200 mmol kg^−1^.

**Fig. 4.**
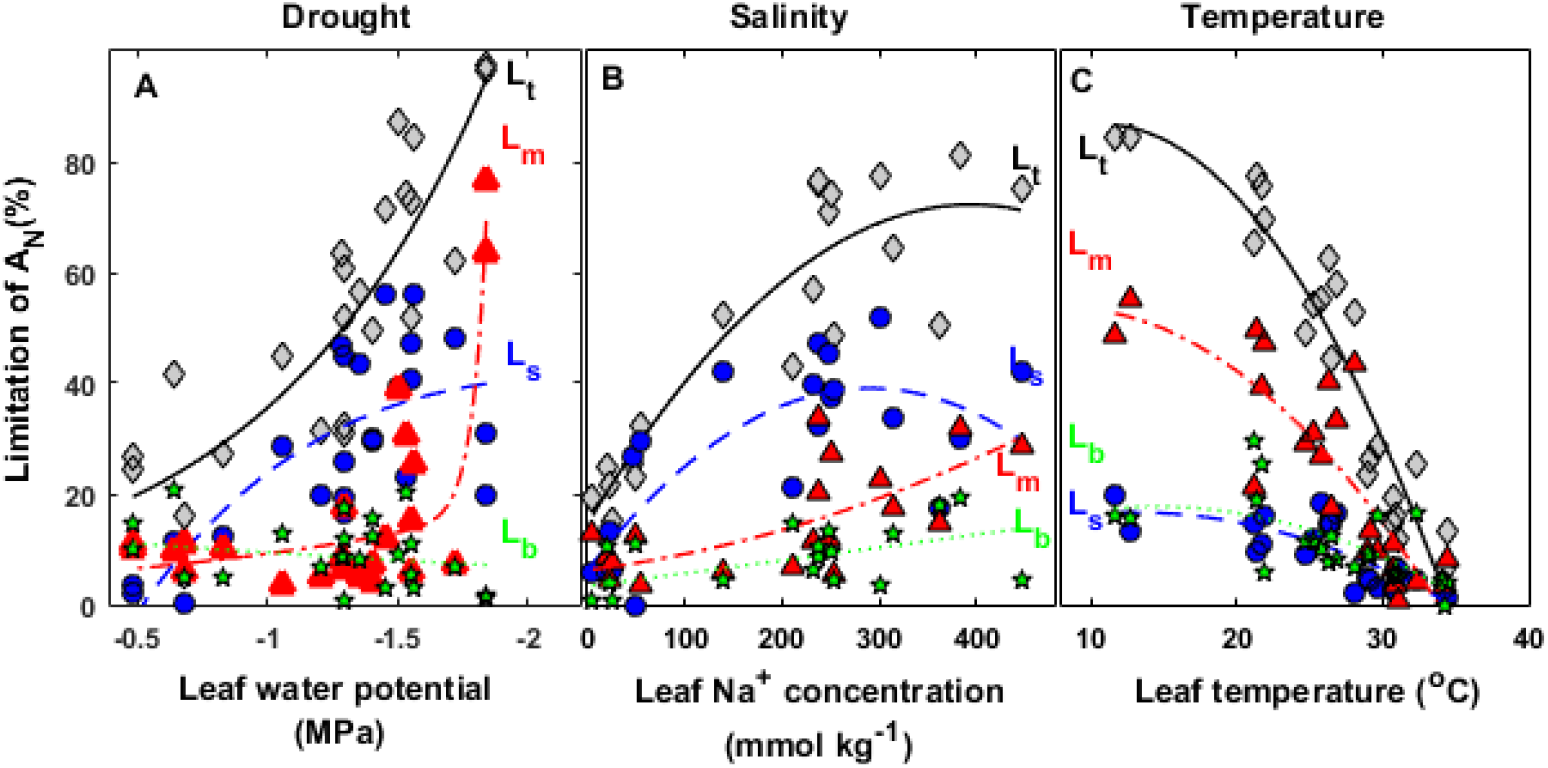
Effects of different environmental stress factors on the overall limitation of photosynthesis (*L_t_*) and its components: stomatal limitation (*L_s_*), mesophyll limitation (*L_m_*) and the biochemical limitation component (*L_b_*). The leaf-level response is expressed differently for each type of stress. **(a)** Drought stress is expressed in terms of leaf water potential, **(b)** salinity is expressed in terms of leaf Na^+^ concentration and **(c)** temperature is expressed in terms of leaf temperature. The continuous line represents the best-fit adjustment for the total limitation (gray diamonds), the line made up of dashes represent the best fit for stomatal limitation (blue circles) and the line made up of dots and dashes represents the best fit for mesophyll limitation (red triangles). The dotted green line represents biochemical limitation of photosynthesis (green stars).

In the temperature experiment (Fig. 4C), all types of limitation increased hyperbolically (*L_t_*: *R*^2^ = 0.89, P<0.0001; *L_s_*: *R*^2^ = 0.6, P<0.0001; *L_m_*. *R*^2^ = 0.72, P<0.0001; *L_b_*: *R*^2^ = 0.36, P=0.0032) during acclimation to decreased leaf temperatures. Unlike acclimation to drought or salinity, acclimation to low temperature (30°C→10°C) was dominated by *L_m_* (∼30-50%), while *L_s_* and *L_b_* (∼10-20%) remained significantly lower.

In order to precisely quantify the stress levels, we created a stress index (*SI*) that expresses a general index for the decrease of leaf physiological performance (excluding GE parameters; Fig. 5). *SI* is composed of the decrease in *K_leaf_, Ψ_P_*, dark-adapted chlorophyll fluorescence (<u>*F_v_*</u>/<u>*F_m_*</u>) and the amount of soluble sugar in the leaf (SC, see Fig. S1). Higher *SI* values indicate stress susceptibility; whereas lower values indicate advanced stress tolerance. SI is correlated with relative growth so that a 50% reduction in growth is equivalent to a *SI* of 0.4 (Fig. 5A). A *SI* of less than 0.5 allows for continued growth, as well as the improved *iWUE* (Fig. 5C) accompanied with increase in the *g_m_*:*g_s_* ratio (Fig. 5B). In addition, it can be seen that SI is significantly correlated with the overall limitation of photosynthesis (*L_t_*) for all treatments combined (*R*^2^ = 0.89, *P* < 0.0001) (Fig. 6). Plotting *L_t_* components (*L_s_, L_m_* and *L_b_*) versus SI revealed that SI is significantly correlated with *L_m_* (*R*^2^ = 0.91, *P* < 0.0001), has a weaker correlation with *L_s_* (*R*^2^ = 0.24, P=0.1) and is not correlated with *L_b_* (*R*^2^ = 0.03, P=0.25).

**Fig. 5.**
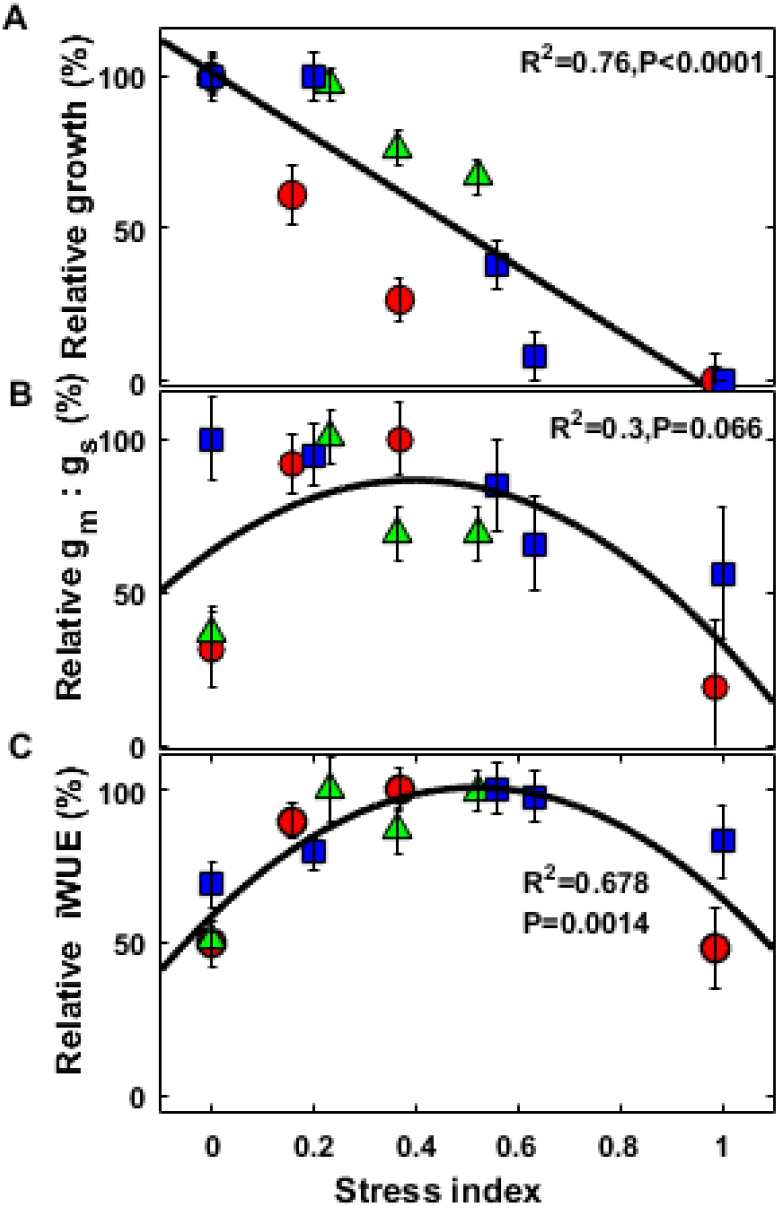
The relationships between **(a)** relative growth, **(b)** relative intrinsic water-use efficiency and **(c)** relative change in the *g_m_*:*g_s_* ratio and the leaf stress index for all of the stress treatments: drought (red circles), salinity (green triangles) and temperature (blue squares). The continuous line represents the best fit for all types of stress combined.

### Correlation of *g_m_* with leaf-water-relations parameters in the different under various stresses

Plotting *g_m_* versus *K_leaf_* shows different patterns for each stress (Fig 7A). In the case of drought stress, *g_m_* and *K_leaf_* were not coordinated. A sharp decline in *K_leaf_* from 9 ± 1.57 to 1.8 ± 0.5 mmol H_2_O m^−2^ s^−1^ MPa^−1^ did not affect *g_m_*. No correlation was found between *K_leaf_* and *g_m_* in the salinity experiment. However, in the temperature experiment, *g_m_* and *K_leaf_* were strongly correlated (*R*^2^ = 0.97, *P* = 0.001). The relationship between *g_m_* and *Ψ_P_* was not correlated with any of the examined types of stress (Fig. 7B). However, we did find a threshold of *Ψ_P_* < 0.4 MPa for the drop in *g_m_* that was observed in the drought-stress experiment.

**Fig. 6.**
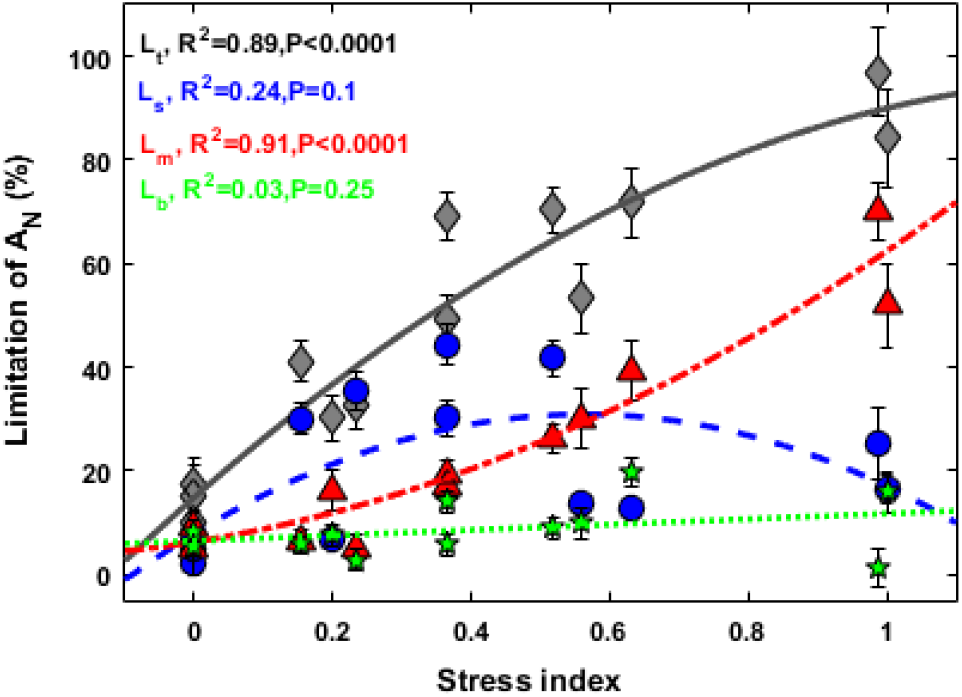
The relationships between the total limitation of photosynthesis (*L_t_* – gray diamonds) and its components: stomatal limitation (*L_s_* – blue circles), mesophyll limitation (*L_m_* – red triangles) and biochemical limitation (*L_b_* – green stars), and the stress index for all of the stress treatments combined. The continuous line represents the best fit for *L_t,_*, the dashed line represent the best fit for *L_s_*, the line made up of dots and dashes represents the best fit for *L_m_* and the dotted line represents the best fit for *L_b_*.

**Fig. 7.**
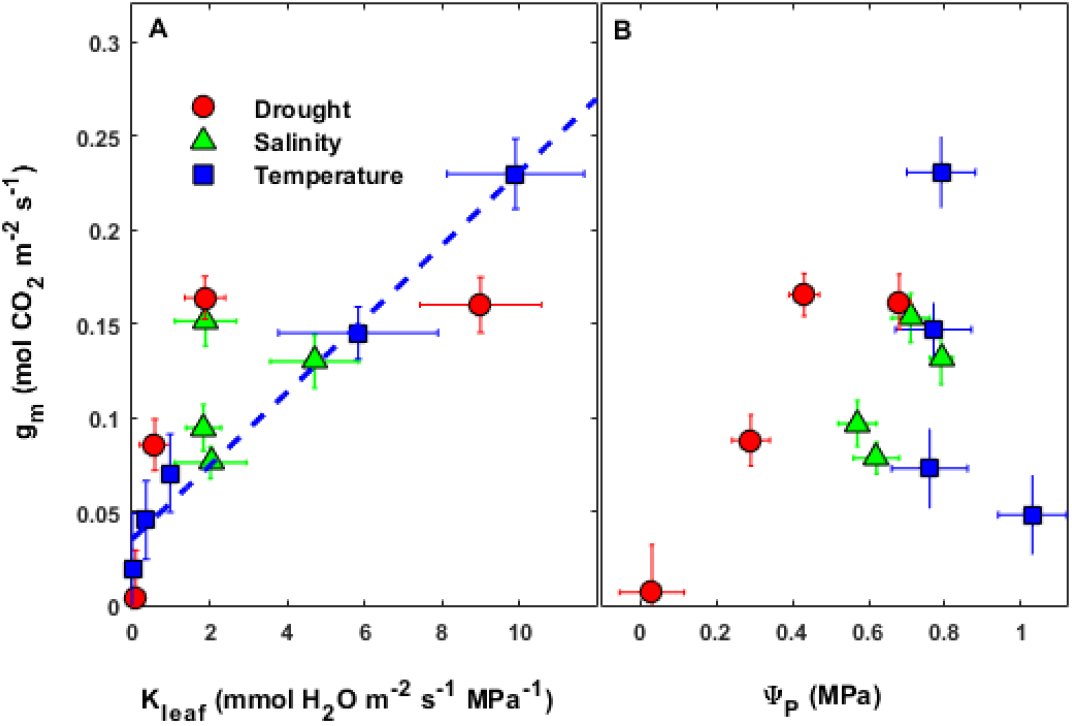
Relationships between mesophyll conductance and **(a)** the estimated leaf hydraulic conductance (*K_leaf_*) and **(b)** the calculated leaf turgor pressure (*Ψ_P_*). The different types of stress are represented as follows: drought (red circles), salinity (green triangles) and temperature (blue squares). The broken blue line represents the best-fit adjustment for the *g_m_*/*K_leaf_* correlation in the temperature experiment.

### Stress, leaf structural traits and *g_m_*

Examination of structural and anatomical leaf characteristics revealed differences between treatments in the drought and temperature experiments (Table 1). In contrast, no significant structural or anatomical changes were found in the salinity experiment. Unlike many other forest trees characterized by a classic dorsoventral mesophyll arrangement, *Ziziphus spina-christi* has a unique mesophyll arrangement that does not include any clear spongy mesophyll (Fig. S3). Acclimation to drought and temperature were each characterized by an increase in leaf mass per area (*LMA*) and leaf width (*L_t_*), accompanied by a decrease in the leaf intercellular spaces (%*IAS*). Analysis of leaf cross-sections revealed a significantly larger mesophyll surface area exposed to the intercellular air space (*S_m_*) and thicker leaves in the low-temperature treatment (16/10°C).

**Table 1.** Leaf structural and anatomical characteristics under long-term drought, salinity and temperature stress. For each treatment, the following information is presented: leaf mass/area (*LMA*), stomatal density (*SD*), leaf width (*L_t_*), mesophyll surface area exposed to intercellular airspaces (*S_m_*) and fraction of intercellular air space (*IAS*). Different letters denote a statistically significant difference between treatment means for each stress factor.

|  | Treatment | LMA<br>(g m<sup>-2</sup>) | SD<br>(n mm<sup>-2</sup>) | L<sub>t</sub><br>(μm) | S<sub>m</sub><br>(m<sup>2</sup> m<sup>-2</sup>) | IAS<br>(%) |
| --- | --- | --- | --- | --- | --- | --- |
| **Drought** | WW | 77.5±2.4 a | 197.9±18 a | 115.7±5.9 a | 9.3±0.2 a | 25.2±1.2 a |
|  | WS1 | 99.7±3.5 b | 200.0±18 a | 116.2±18.4 a | 8.9±0.1 a | 15.3±1.7 b |
|  | WS2 | 124.5±3.5 c | 196.0±31 a | 146.4±15.2 a | 10.2±0.2 b | 14.9±1.0 b |
|  | *P* value | 0.0003 | n.s | n.s | 0.014 | 0.0065 |
| **Salinity**<br>**(mM)** | 0 | 108.7±3.7 a | 229.3±13.0 a | 134.0±6.0 a | 8.8±0.4 a | 23.8±0.4 a |
|  | 30 | 91.5±11.6 a | 227.5±25.5 a | 122.5±11.4 a | 9.8±1.6 a | 21.5±0.8 a |
|  | 60 | 120.6±11.4 a | 219.0±32.6 a | 142.4±7.6 a | 10.4±0.5 a | 16.7±1.1 a |
|  | 90 | 106.7±3.4 a | 196.9±19.4 a | 145.4±14.4 a | 10.6±1.1 a | 21.0±1.2 a |
|  | *P* value | n.s | n.s | n.s | n.s | n.s |
| **Temperature**<br>**(°C day/night)** | 34/28 | 77.8±4.3 a | 319.4±26.0 a | 177.4±5.7 ab | 8.4±0.7 a | 21.5±2.2 a |
|  | 28/22 | 75.1±4.5 a | 274.8±26.9 ab | 174.4±5.9 ab | 7.3±0.9 a | 21.7±2.3 a |
|  | 22/16 | 87.7±4.0 ab | 252.2±24.4 ab | 160.6±5.4 b | 6.9±0.9 a | 12.8±2.0 b |
|  | 16/10 | 93.9±5.5 b | 197.7±32.8 b | 196.8±7.2 a | 11.3±0.2 b | 14.6±2.8 ab |
|  | *P* value | 0.0239 | 0.0667 | <.0001 | 0.0007 | 0.0361 |

When data from all of the stress treatments were pooled, none of the examined structural or anatomical traits exhibited any correlation with *gm* (Fig. 8). When the data sets collected for the different experiments were examined separately, we found a tight negative correlation between *g_m_* and *LMA* in the drought experiment (*R*^2^ = 0.52, *P* = 0.26) and in the temperature experiment (*R*^2^ = 0.17, *P* = 0.037) (Fig. 8A). *g_m_* was correlated with *L_t_* only in the drought experiment (*R*^2^ = 0.79, *P* < 0.01) (Fig. 8B). Among the pooled data, no relationship was observed between *g_m_* and *S_m_* (Fig. 8C). A correlation between *g_m_* and the %*IAS* was observed only in the temperature experiment (*R*^2^ = 0.34, *P* < 0.01) (Fig. 8D).

**Fig. 8.**
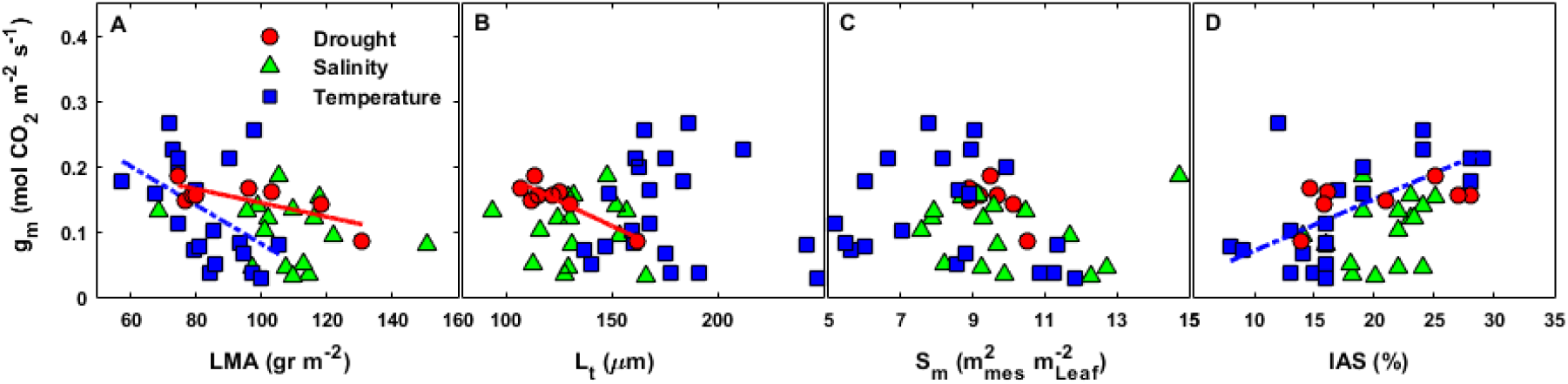
Relationships between *g_m_* and leaf structural and anatomical traits under different types of environmental stress: drought (red circles), salinity (green triangles) and temperature (blue squares). **(a)** Leaf mass per area, **(b)** leaf thickness, **(c)** mesophyll surface area exposed to the intercellular airspace (*S_m_*) and **(d)** percentage of leaf intercellular airspace (*IAS*).

## Discussion

Our method of precisely defining stress allowed us to compare three important types of abiotic stress and investigate the physiological acclimation of our species of interest to different levels of those types of stress (Figs. 1,5,6). For all of our experiments, we examined the youngest leaves, which developed under the applied stress conditions. In this manner, we were able to avoid the well-known negative effects of leaf senescence on photosynthesis-limiting factors (Niinemets *et al*., 2005). As expected, growth and photosynthesis gradually declined during the course of acclimation to each of the different types of stress (Figs. 1,2). Quantitative-limitations analysis demonstrated that for all types of stress, photosynthesis is mostly limited by the diffusion of CO_2_ into the chloroplasts, through *gsc* and *gm* (∼90% for drought and salinity, ∼80% temperature); whereas biochemistry has only a minor effect (∼10% drought and salinity, ∼20% temperature; Fig. 4). These findings are consistent with earlier reports on trees’ acclimation to drought and salinity (Bongi and Loreto, 1989; Grassi and Magnani, 2005; Galmés *et al*., 2007a; Perez-Martin *et al*., 2009; Keenan *et al*., 2010; Limousin *et al*., 2010; Egea *et al*., 2011; Cano *et al*., 2013, 2014). As for growth temperature, information about long-term acclimation to higher temperatures is currently available for only one tree species *(Eucalyptus regnans;* Warren, 2008b). However, that report did not involve a quantitative analysis of photosynthetic limitations that included *L_b_*.

Though *g_sc_* and *g_m_* were the main limiting factors for all of the examined types of stress, their relative contributions to the diffusive limitation differed between the types of stress (Figs. 3,4). The limitation of photosynthesis during acclimation to drought and salinity was controlled mostly by stomata (*L_s_*), whereas mesophyll limitation (*L_m_*) was important only under progressive stress (*LWP* >1.5 MPa or leaf Na^+^ > 200 mmol kg^−1^). However, the limitation of photosynthesis during acclimation to lower growth temperatures (34→10°C) was significantly controlled by the mesophyll (∼30-50%), while the stomata and biochemical limitation accounted for less than 20% of the observed reduction in the rate of photosynthesis.

It is well established that in forest trees a reduction in *g_s_* is the earliest response to drought and salinity, as well as the most perceptible photosynthesis limitation (Ziska *et al*., 1990; Wilson *et al*., 2000; Maseyk *et al*., 2008; Limousin *et al*., 2010). Stomatal closure is the most well-described characteristic of leaf stress avoidance (Raschke, 1975; Levitt, 1980). Unlike *g_s_*, which is actively regulated, *g_m_* does not have any known regulatory mechanism (Parkhurst, 1994; Tholen *et al*., 2012; Barbour, 2017). Mesophyll cells are protected from damage caused by dehydration as they are partially hydraulically buffered from the transpiration stream (Buckley *et al*., 2017). Most studies indicate that *g_m_* drops at a later stage of drought stress, but the initial influence and magnitude varies greatly among species (Galmés *et al*., 2007a; Cano *et al*., 2014; Théroux-Rancourt *et al*., 2014; Zhou *et al*., 2015). Stress acclimation of *g_m_* and its role in the limitation of photosynthesis are relatively difficult to predict. Our proposed stress index may explain trees’ vast investment in mesophyll protection. As the stress index (*SI*) increases, there is a linear decrease in the growth rate (Fig. 5A). In parallel, iWUE and *g_m_*:*g_s_* increase until *SI* reaches 0.4 (Fig. 5 B,C). This is still the avoidance stage, in which the leaf is effectively acclimated to its new non-optimal conditions. Later on (when *SI* exceeds 0.4), the ratio of *g_m_* to *g_s_* is impaired, which may represent the breakdown of the *g_m_* protective barriers (i.e., stomatal regulation and hydraulic compartmentalization). We suggest that *g_m_* drops only at the stage at which stress prevails (i.e., when the stomatal protective mechanism is less effective). Moreover, the integrative relationship for photosynthesis limitation accounts for all types of stress, revealing a strong correlation with *SI* only for *L_m_* (Fig. 6). This also emphasizes how *g_m_* is a key factor in tolerance and acclimation to non-optimal conditions.

It has been suggested that the exact point at which *g_m_* begins to decrease is a determining factor for drought resistance across different species and genotypes (Cano *et al*., 2014; Théroux-Rancourt *et al*., 2014, 2015; Barbour & Kaiser, 2016). Our results are in line with those previous findings and further suggest that a stable low *L_m_* may be a strong indicator of stress tolerance in acclimation to salinity and low temperature.

Our results also showed that improved *iWUE* under intensified drought and salinity stresses was made possible by a greater decline in *g_s_*, rather than *g_m_* (a high *g_m_*:*g_s_* ratio; Fig. 3 A,B,C,D). On the other hand, relatively low *iWUE* at low temperatures was accompanied by a low *g_m_*:*g_s_* ratio, mostly due to the opposite effect: a stronger influence of *g_m_* relative to *g_s_*; Fig. 3E,F). It was recently suggested that trees originating in a xeric environment have higher *g_m_*:*g_s_* ratios than mesic trees (Stpaul *et al*., 2012; Cano *et al*., 2014; Zhou *et al*., 2014; Peguero-Pina *et al*., 2017). It has also been proposed that species from warmer climates exhibit a steep *g_m_*-temperature response (leading to low *g_m_*:*g_s_*); whereas the *g_m_* levels of species from cooler climates are slightly change in response to changes in temperature (von Caemmerer & Evans, 2015). Our results are congruent with these two hypotheses.

Under stress conditions, leaves exhibited structural rearrangements, including an increase in LMA and reduced IAS (Table 1), which may have a negative influence on *g_m_* (Poorter *et al*., 2009; Tomás *et al*., 2013; Muir *et al*., 2014; Onoda *et al*., 2017). However, increases in LMA were accompanied by increases in *S_m_*, which is a major positive determinant of *g_m_* under optimal conditions (von Caemmerer and Evans, 1991; Peguero-Pina *et al*., 2017). In *Ziziphus*, it appears that mesophyll anatomy alone does not explain *g_m_* acclimation to drought, salinity or temperature stress (Fig. 8).

The fact that *g_m_*, similarly to *g_s_*, can change rapidly within seconds or minutes in response to environmental or internal stimuli, indicating that leaf anatomy is not the only determining factor (Flexas *et al*., 2007). There is some evidence that CO_2_ and water can both pass through aquaporins (Terashima and Ono, 2002; Flexas *et al*., 2006b). Thus, *g_m_* and *K_leaf_* are expected to be positively correlated under optimal conditions (Flexas *et al*., 2013). However, it is still unclear whether this correlation holds under stress conditions (Loucos *et al*., 2017).

In our case, a correlation between *g_m_* and *K_leaf_* was observed only in the temperature experiment (Fig. 7A). This finding strengthens the argument that shared pathways for CO_2_ and water lead to coordination between *g_m_* and *K_leaf_* and that that process is temperature dependent (Bernacchi *et al*., 2002). Although we observed a sharp drop in *K_leaf_* in the drought experiment, under those conditions, *g_m_* decreased only slightly. This suggests that mesophyll cell-to-cell CO_2_ pathways are not influenced by moderate drought and that only severe drought will cause a coupled decline of both *K_leaf_* and *g_m_*. Even so, the relationship between *g_m_* and *K_leaf_* may vary among species and may also be related to anatomical properties of the mesophyll (Xiong *et al*., 2016, 2017). Further study of these relationships under different stress conditions is needed.

We also demonstrated that *g_m_* was not correlated with *Ψ_P_* for any of the examined types of stress (Fig. 7B). *g_m_* appears to respond to drought at *Ψ_P_* levels lower than 0.4 MPa. Still, threshold-like responses and mechanisms remain obscure. We suggest that *g_m_* becomes severally impaired only when the mesophyll becomes flaccid. Flaccid mesophyll tissues may go through transformative changes involving cell collapse and reduced IAS (Levitt and Ben Zaken, 1975). This would reduce leaf internal effective CO_2_ diffusion pathways and explain low *g_m_* (von Caemmerer and Evans, 1991). Other possible explanations are based on the idea that deactivation of aquaporins occurs only under conditions of dramatic turgor loss (Miyazawa *et al*., 2008). However, there is currently no known mechanism for aquaporin-turgor sensing in the mesophyll cells and further research is needed to explain this phenomenon.

For salinity stress, the point at which *g_m_* dropped can be explained in other ways. Plants responses to salt stress are generally accepted to have two components: an osmotic component and an ionic component (Munns & Tester, 2008). The primary osmotic stress characterized by a reduction in both the water potential and the osmotic potential results in a rapid decrease in *g_s_*. Following that rapid effect, under prolonged salt stress, ion imbalance and toxic levels of Na^+^ and Cl^−^ inhibit enzyme activity and protein synthesis (Chaves *et al*., 2009; Shapira *et al*., 2009). Yet, it is still unclear whether the reduction in *g_m_* can be attributed to the osmotic effect or to the ionic toxicity (Delfine *et al*., 1999; Centritto *et al*., 2003; Chen *et al*., 2015; Razzaghi *et al*., 2015). In our case, the lowest *LWP* measured for the extreme salinity treatment (90 mM, leaf Na^+^ > 300 mmol kg^−1^) was −1.1 MPa and, at that level, the leaves remained turgid (Fig. 7B). This indicates that the leaves were osmotically acclimated and that the reduction in *g_m_* was triggered mostly by leaf accumulation of Na+ and Cl^−^ (i.e., ionic factors).

## Conclusions

In this work, we have compared for the first time the photosynthesis limitation resulting from acclimation to three different types of stress. Such a comparison can lead to a better understanding of which aspects of photosynthesis limitation are dominant under each type of stress. It may also help to predict how this species adapts to its environment and to future climate change. Our results clarify that *Z. spina-christi* is unable to withstand low temperatures due to an extremely sensitive *g_m_*. On the other hand, this species exhibits high tolerance to drought and salinity characterized by a low *g_s_* accompanied by a stable and constant *g_m_*. With ongoing climate changes, the Mediterranean basin is likely to experience more frequent and intense drought conditions, accompanied by higher winter temperatures (Alpert *et al*., 2008). These trends may give *Z. spina-christi* an ecological advantage that may lead to its becoming more widely distributed across this region in future decades. Consequently, this tree may serve as an indicator species for climate change, which is pushing the limits of the distribution ranges of many species.

## Acknowledgment

This project was supported by a grant (No. 10-01-005-13) obtained from the Jewish National Fund (JNF) for the study of the ecophysiology of *Ziziphus spina-christi* under changing climate conditions.

## Author Contributions

YZ – planning and performing the experiments, data analysis, writing, IS-performing anatomy experiments, AS – planning the experiments and writing.

## Suplementary Data

**Table S1.** Parameters derived from the analysis of the A_N_/C_c_ and A/Q curves for the different types of environmental stress (drought, salinity and temperature). Laisk-method estimation of the mitochondrial respiration in the light (Rd) and the apparent intercellular CO_2_ photo-compensation point (C_i_*). Valentini method estimation of the total electron-transport rate (J_t_) calculated using the, electron-transport rate derived for carboxylation (J_c_), electron-transport rate derived for oxygenation (J_o_), Jc:Jo ratio and CO_2_ emitted in photorespiration (F). Different letters denote a statistically significant difference between the treatments means for each type of stress.

|  | Treatment | "Laik method " |  |  | "Valentiny method " |  |  |  |
| --- | --- | --- | --- | --- | --- | --- | --- | --- |
| | | $R_d$<br>($\mu mol CO_2 m^{-2} s^{-1}$) | $C_i^*$<br>($\mu mol mol^{-1}$) | $J_t$<br>($\mu mol e^- m^{-2} s^{-1}$) | $J_c$<br>($\mu mol e^- m^{-2} s^{-1}$) | $J_o$<br>($\mu mol e^- m^{-2} s^{-1}$) | $J_c:J_o$ | $F$<br>($\mu mol CO_2 m^{-2} s^{-1}$) |
| Drought | WW (n = 7) | 1.0 ± 0.08 a | 38.4 ± 2.0 a | 155.1 ± 5.9 a | 114.0 ± 3.0 a | 41.1 ± 3.0 a | 3.0 ± 0.12 a | 5.1 ± 0.4 a |
|  | MWS1 (n = 7) | 0.82 ± 0.08 b | 36 ± 1.3 a | 132.0 ± 7.9 b | 84.7 ± 4.1 b | 47.3 ± 4.3 a | 1.9 ± 0.16 b | 5.9 ± 0.5 a |
|  | MWS2 (n = 7) | 0.7 ± 0.08 b | 36.4 ± 1.5 a | 99.4 ± 6.6 c | 54.8 ± 3.4 c | 44.6 ± 3.6 a | 1.3 ± 0.13 c | 5.5 ± 0.4 a |
|  | SWS (n = 7) | 0.6 ± 0.1 b | 39.0 ± 1.5 a | 81.7 ± 5.7 d | 34.6 ± 3.0 d | 47.1 ± 3.1 a | 0.75 ± 0.1 d | 5.8 ± 0.3 a |
|  | P value | 0.0012 | n.s | <0.0001 | <0.0001 | n.s | <0.0001 | n.s |
| Salinity | 0 mM (n = 6) | 0.95 ± 0.1 a | 34.1 ± 1.5 a | 159.7 ± 6.6 a | 107.4 ± 4.6 a | 52.3 ± 2.7 a | 2.0 ± 0.08 a | 6.5 ± 0.3 a |
|  | 30 mM (n = 5) | 0.93 ± 0.04 a | 34.0 ± 1.9 a | 148.9 ± 8.0 a | 98.2 ± 5.6 a | 49.7 ± 3.3 a | 2.0 ± 0.1 a | 6.2 ± 0.4 a |
|  | 60 mM (n = 6) | 0.84 ± 0.05 a | 36.9 ± 1.4 a | 105.7 ± 7.0 b | 63.9 ± 4.7 b | 42.2 ± 2.8 a | 1.5 ± 0.09 b | 5.2 ± 0.36 a |
|  | 90 mM (n = 6) | 0.83 ± 0.04 a | 32.1 ± 1.3 a | 111.5 ± 7.4 b | 59.9 ± 5.2 b | 51.7 ± 3.0 a | 1.1 ± 0.09 b | 6.4 ± 0.38 a |
|  | P value | n.s | n.s | <0.0001 | <0.0001 | n.s | <0.0001 | n.s |
| Temperature | 34°C (n = 6) | 1.2 ± 0.13 a | 41.8 ± 2.9 a | 178.8 ± 9.1 a | 122.0 ± 5.0 a | 56.8 ± 4.0 a | 2.1 ± 0.2 a | 7.1 ± 0.5 a |
|  | 28°C (n = 6) | 0.9 ± 0.1 b | 39.0 ± 2.2 a | 140.3 ± 8.3 b | 104.1 ± 4.7 a | 36.1 ± 3.7 b | 2.8 ± 0.18 ab | 4.5 ± 0.4 b |
|  | 22°C (n = 6) | 0.6 ± 0.1 c | 29.9 ± 2.2 b | 103.1 ± 7.7 c | 70.6 ± 4.7 b | 32.4 ± 3.7 b | 2.2 ± 0.18 ab | 4.0 ± 0.4 b |
|  | 16°C (n = 6) | 0.4 ± 0.12 c | 28.6 ± 2.0 b | 54.5 ± 7.7 d | 33.9 ± 5.0 c | 20.5 ± 4.0 bc | 1.8 ± 0.19 bc | 2.5 ± 0.5 bc |
|  | 10°C (n = 5) |  |  | 26.7 ± 9.1 d | 14.0 ± 5.5 c | 12.7 ± 4.4 c | 1.1 ± 0.20 c | 1.5 ± 0.5 c |
|  | P value | <0.0001 | 0.001 | <0.0001 | <0.0001 | <0.0001 | <0.0001 | <0.0001 |

**Fig. S1** Changes in soluble carbohydrate (SC) and starch content in the **(a)** drought, **(b)** salinity and **(c)** temperature treatments.

**Fig. S2** Correlation of mesophyll conductance (*g_m_*) estimated using the methods described by Harley *et al*. (1992) and Ethier and Livingston (2004) for the drought (red circles), salinity (green triangles) and temperature treatments (blue squares).

**Fig. S3** Representative samples of leaf anatomical cross-sections for the control and extreme treatment for each type of stress, showing differences in mesophyll shape and density.

